# ECM-body: A cell-free 3D biomimetic scaffold derived from intact planarian body

**DOI:** 10.1101/763714

**Authors:** Ekasit Sonpho, Chanida Wootthichairangsan, Miyuki Ishida, Takeshi Inoue, Kiyokazu Agata, Anchuleerat Maleehuan, Komgrid Charngkaew, Nusara Chomanee, Saengduen Moonsom, Patompon Wongtrakoongate, Arthit Chairoungdua, Puey Ounjai

**Affiliations:** Department of Biology, Faculty of Science, Mahidol University, Thailand 10400; Center of Excellence on Environmental Health and Toxicology (EHT), Office of Higher Education Commission, Ministry of Education, Thailand, 10400; Department of Life Science, Faculty of Science, Gakushuin University, Tokyo, Japan 171-8588; Department of Pathology, Faculty of Medicine Siriraj Hospital, Mahidol University, Thailand 10700; Department of Protozoology, Faculty of Tropical Medicine, Mahidol University, Thailand 10400; Department of Biochemistry, Faculty of Science, Mahidol University, Thailand 10400; Department of Physiology, Faculty of Science, Mahidol University, Thailand 10400

**Keywords:** Extracellular matrix, Planarian, Decellularization, Scaffold, Tissue engineering, ECM-body

## Abstract

Extracellular matrix (ECM) plays key roles in shaping fates of stem cells, not only by providing suitable niche but also by mediating physical and biochemical cues. Despite intensive investigations on regeneration, the roles of ECM on fate determination of stem cells in animal with great regenerative potency, such as planarian, remained unclear. Here, we developed a method to decellularizing and isolating extracellular matrix from planarians. Although the isolated scaffold appears translucent, it contains all the internal features, resembling the structure of intact planarian, and which we thus called “ECM-body”. Nuclear staining demonstrated that ECM-body contains very little or no cell remained. Histological sections displayed a well-preserved morphological integrity of the specimen. Scanning electron microscope showed porous surface on ECM-body, potentially suitable for housing cells. Furthermore, our preliminary experiment suggested that ECM-body can be utilized as biomimetic scaffold for cell culture as it may support survival of injected neoblasts.

## Introduction

Planarians have been widely utilized for studying regeneration as they are able to regenerate their body parts within weeks (1–3). Using planarians as models, various processes responsible for regeneration and development have been thoroughly elucidated (3–6). Furthermore, the detailed characterization of stem cell types as well as genes involved in tissues homeostasis and organ formation in planarians have also been established (7–9). It has been shown that the stemness of adult stem cells require physical interactions between the cells and the ECM (10). ECM plays key roles in modulating the dynamic microenvironment by providing suitable physical habitat for the resident cells, allowing the cell to develop and differentiate into specialized cells and governing cell to cell communications through controlled distribution of chemicals and biochemical signals (11). Therefore, various attempts have been made to create suitable biomimetic 3D scaffold in order to simulate the microenvironment of ECM, especially for understanding the complex interaction between the ECM and the stem cells (12). Indeed, a method of choice to prepare cell-free ECM is so-called decellularization due to the fact that the ECM obtained from this method could potentially provide suitable microenvironment for the cells, offering not only mechanical and physical cues but also some cryptic biochemical signals that are involved in regulating growth and developmental processes (13).

To date, various protocols have been developed for decellularizing various mammalian tissue systems (14–16). However, there was only limited information regarding isolation of ECM of invertebrates (17,18). To our knowledge, there is currently no decellularization protocol developed for planarian tissues.

Here, we established a protocol for preparing intact decellularized ECM, which we called ECM-body, from freshwater planarians, *Dugesia* sp. (Thai isolate) and *Dugesia japonica* and preliminarily explored the possibility to utilize ECM-body as biomimetic scaffold for 3D cultivation of planarian neoblasts. Our work offered broad application in investigating the role of ECM components not only in regeneration but also in stem cell biology.

## Results and Discussions

### Optimization of Decellularization Procedure for Freshwater Planarians

Decellularization is a technique to prepare three-dimensional biomimetic scaffold for tissue engineering and microenvironmental studies (13–16). The goal of successful decellularization is to completely get rid of all the cellular components while faithfully preserving the intact architecture of ECM as well as other potential microenvironmental factors. Although various methods have been established for decellularizing mammalian organs (12), the decellularization of freshwater planarians was obviously quite challenging because their bodies are often quickly disintegrated upon the death of animal. Due to the rapid growth and the ease of cultivation, we chose to initially optimized a protocol for whole-body decellularization using *Dugesia* sp. (Thai isolate). Sodium dodecyl sulfate (SDS) has been utilized as an effective agent for removing nuclei from decellularized tissues (19), we therefore selected SDS as the first line of decellularizing agent. Although the resulting ECM-body prepared in this condition appeared to largely resemble the shape of planarians (Fig 1A and B), the internal organization of ECM-body prepared using this method appeared to be less defined. To better preserve the internal and external 3D organization of planarian, we decided to include a step to stabilize the planarian body using cross-linking agents such as formaldehyde prior to the decellularization process. The agent not only help stabilizing complex internal structure of the worm but also quickly terminate the animal, preventing cellular apoptosis which might possibly affect the integrity of resulting ECM (20). The ECM-body prepared by this improved procedure appeared translucent with well-preserved internal organ systems of the intact planarians i.e. digestive tract, eye spots and pharynx (Fig 1A and C). Due to the fact that the stabilization could potentially cause negative impact on the efficiency of cellular removal, we rescreened the panel of detergents, consisting of Triton X-100, Tween-20 and SDS, to seek for the most effective decellularizing agent for removing the cell from stabilized planarian bodies. Although all the selected decellularizing agents were able to successfully dissolve cellular content from the stabilized worms, cellular removal capability of SDS-based extraction was still superior in comparison with the other two selected detergents (Fig 1D). It should be noted that the internal structure of digestive tract of the worm within 1 hour after treatment with low concentration of SDS (Fig 1D). Translucent ECM-body containing clear intact internal structures was achieved within 4 hours (Fig 1D). To verify the cellular removal efficacy of different detergents, the amount of residual DNA in the sample was determined at various time points during the treatment (Fig 1E). The results confirmed that the decrease of residual DNA 0.08% SDS was significant faster than the other two detergents tested (Fig 1E). Therefore, SDS was chosen as a detergent of choice for subsequent decellularization experiment.

**Fig. 1.**
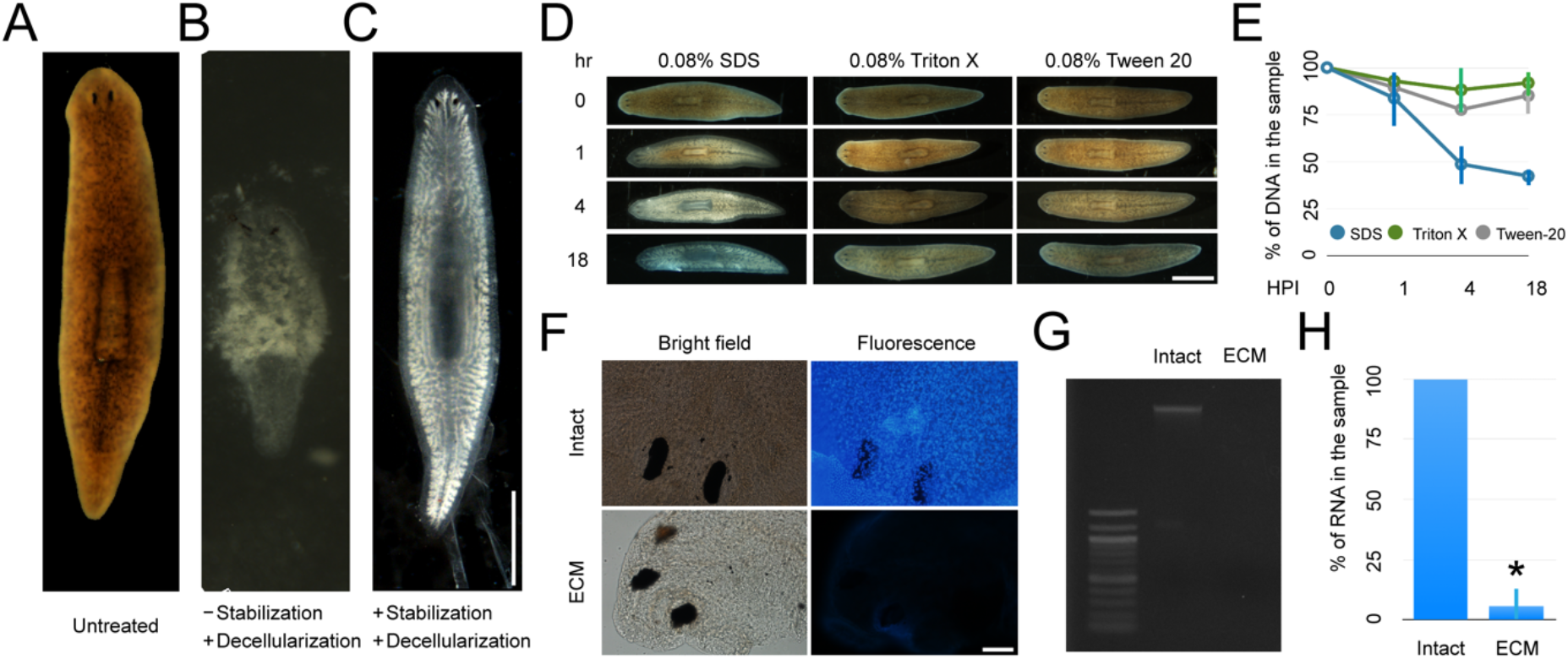
Optimization of planarian ECM-body Isolation. (A) The morphology of untreated planarians, (B) decellularization without and (C) with stabilization. Scale bars are 1 mm (D) The decellularization efficiency among different detergents. Scale bars are 1 mm (E) The comparison of DNA content left in planarians after decellularizing with different detergents. Error bar indicated standard deviation from 2 independent experiments, with 10 planarians per experiment, ANOVA, (P<0.05). (F) DAPI stained planarian and ECM-body. Scale bars are 50 μm (G) Gel electrophoresis for gDNA (H) The amount of RNA was found in the ECM-body. The data was from 3 independent experiments, with 10 planarians per experiment, ANOVA, (P<0.05).

### Characterization of Acellular Planarian ECM-body

As one of the minimal criteria for acceptable decellularization is the lacking of visible nuclear material, the ECM-body was stained with 4’, 6-diamidino-2-phenylindole (DAPI) (21). The results demonstrated that there was no nucleus remained in the isolated ECM (Fig 1F). However, weak signals of DAPI can still be observed in the background perhaps from the residual nucleic acid fragment remained attached to the scaffold. However, agarose gel electrophoresis revealed no band of remaining genomic DNA (Fig 1G). Further fluorometric quantification of RNA also displayed marked decrease in the RNA content in the ECM body (Fig 1H).

To verify the integrity of the ECM architecture, ECM body was stained with Masson’s trichrome dyes (22). Histological cross sections at the pharyngeal plane elucidated intact organization of fibrous ECM structures similar to that of the untreated worm (Fig 2 A and B). Notably, little or no cell were found remained in the scaffold. Using scanning electron microscopy (SEM), the surface topography of intact worm and isolated ECM-body were characterized. The results displayed that the dorsal and ventral sides of the planarian body were easily distinguishable as the former are covered with mucus layer while the latter contains tremendous ciliature on the surface (Fig 2C and D). On the contrary, both dorsal and ventral surfaces of the ECM body displayed similar porous structure. This is in great agreement with the corresponding three-dimensional fibrous network observed in the histological staining (Fig 2E). It should be noted that the ECM-body visualized using SEM appeared flattened. This effect was likely caused by the dehydration during SEM sample preparation. The flattening of the ECM-body may also suggest that ECM body comprised soft and airy 3D scaffold. Taken together, this thus confirmed that our developed protocol did not only effectively remove residential cells from the worms but also provide faithful preservation of intact internal architecture of cell free matrix. Our optimized decellularization protocol is summarized in Fig 3A.

**Fig. 2.**
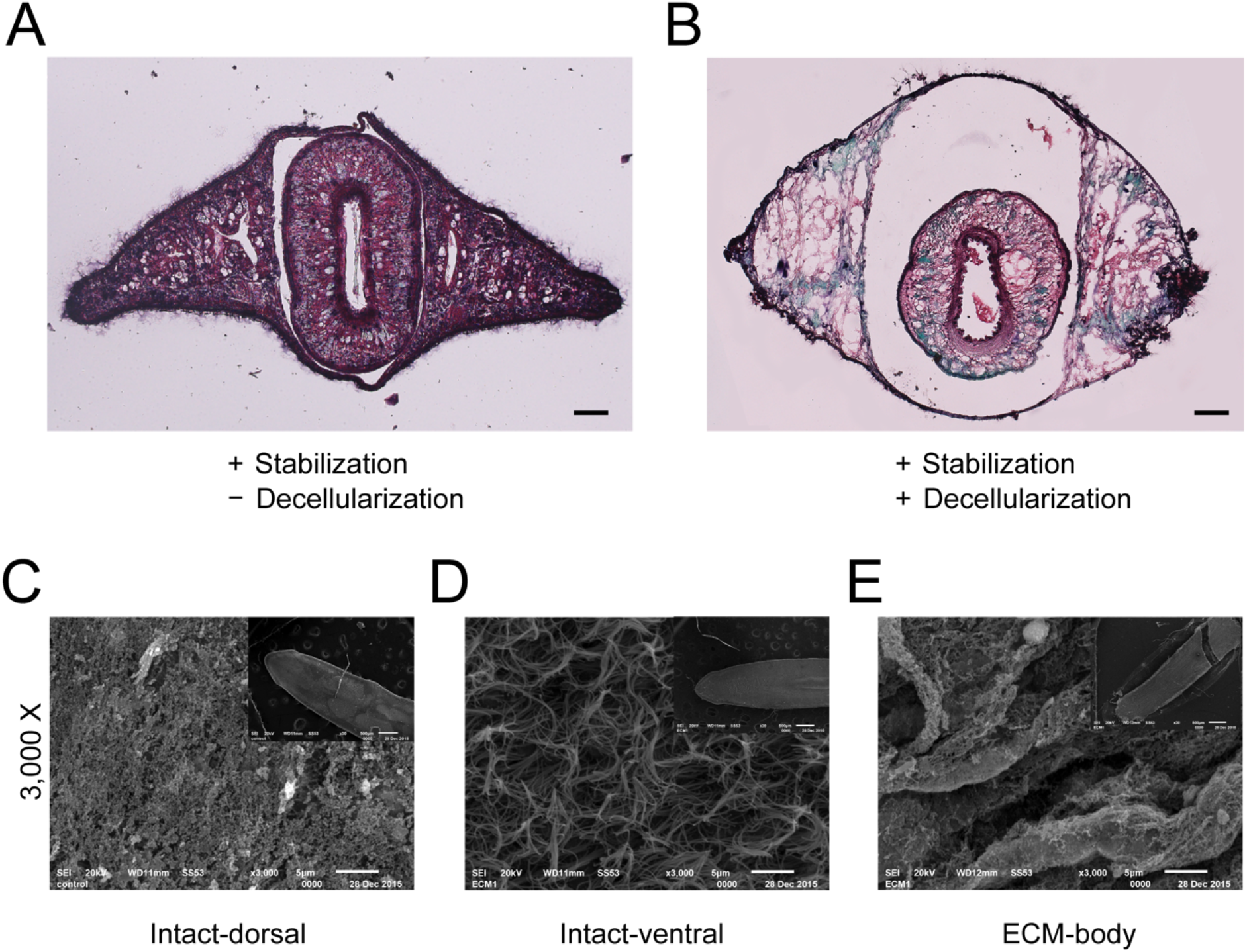
Physical assessment of ECM-body. Histological cross sections at the pharyngeal plane of the planarian (A) and ECM-body (B) stained with Masson’s trichrome dyes. Scale bars are 50 μm. (C) External surface of dorsal side of intact planarian (D) ventral side of intact planarian (E) ECM-body. The insect illustrates the expand view of samples. Scale bars in the insect is 500 μm and scale bar in large image is 10 μm

**Fig. 3.**
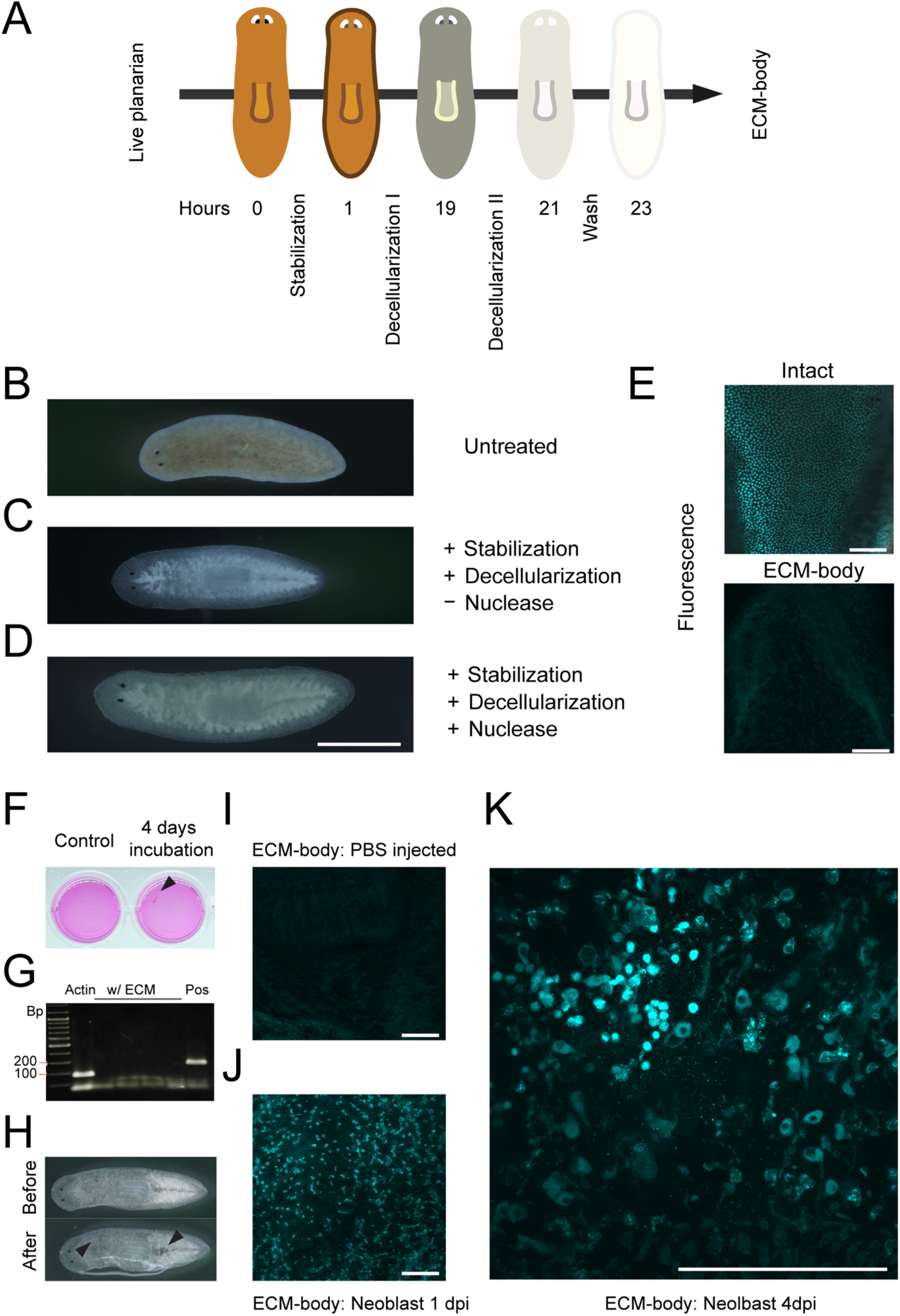
Compatibility of decellularization procedure with *Dugesia japonica*. (A) The diagram showed the overview of the optimized protocol for decellularization of Thai isolates planarian. The ECM of *D. japonica* were isolated based on the procedure optimized in Thai planarians. (B) Untreated planarians, (C) stabilized-decellularization without and (D) with nuclease. Scale bars are 1 mm (E) DAPI stained planarian and ECM-body. Scale bars are 100 μm. (F) The clarity of cell culture media after 4 days of incubation with ECM-body. Arrowhead indicated the ECM-body (G) PCR for mycoplasma detection from co-culture of ECM-body and NTERA2 cell line. (H) The appearance of ECM-body before and after microinjection. Arrowheads indicated the site of cell injection. (I) DAPI staining of the PBS injected inside ECM-body. Scale bar is 100 μm. (J) DAPI staining of the injected cells cultured inside ECM-body for 1 days. Scale bar is 100 μm. (K) Hoechst 33342 staining of the microinjected cells cultured inside ECM-body for 4 days. Scale bar is 100 μm.

### Preliminary Utilization of ECM-body as Three-Dimensional Scaffold

To explore the potential of utilizing the ECM body as a biological 3D scaffold for replanting of the neoblasts, we shifted our attention to a well-established helminth model, *Dugesia japonica* Due to the availability of genomic information and an established protocol for FACS sorting of neoblast (23). Without any modification, the intact ECM-body from *D. japonica* can be similarly prepared using our established procedure (Fig 3B-C, E). To further reduce the possible influence from nucleic acid remnants on the behaviour of the replanted cells (24), the ECM body was further subjected to DNase and RNase treatment. We observed that the additional nuclease treatment did not affect overall morphology of the ECM-body (Fig 3C-D).

Although the use of formaldehyde in stabilization solution should kill all the potential contaminating microbes (25), to ensure the suitability of exploitation of ECM-body as a 3D scaffold in cell culture, the ECM-body was further assayed for bacterial and fungal contamination by incubating in mammalian cell culture media. As a result, based on the colour and clarity of the media, no sign of microbial growth was observed after 4 days of incubation (Fig 3F). Moreover, a polymerase chain reaction (PCR) using specific to mycoplasma validated that the ECM-body prepared with our method was mycoplasma free (Fig 3G).

Finally, in order to set stage for future utilization of the ECM-body in advancing the study of interaction between planarian stem cell and niche, we performed *ex vivo* recellularization of the FACS isolated X1 fraction (FACS sorted S or G2/M phase neoblasts) from planarians into sterile ECM-body both at the anterior and posterior using microinjection (26). Upon injection, some swelling effects were observed at the sites of microinjection, possibly due to the pressure build up during the microinjection (Fig 3H). After incubation at 24 °C, the microinjected ECM-body were taken out at 1 and 4 post-injection (dpi) for evaluating the cell retention. Of interest, the Hoechst stain revealed random distribution of cell and cell clusters in the ECM, suggesting that ECM-body was unlikely to contain residual fixatives and was not toxic to the replanted stem cells. In fact, the ECM-body might be able to support the settlement of neoblast as it could provide the site for cell attachment (Fig. 3I-K). It should be noted that the viability of the seeded the X1 neoblasts in ECM-body was maintained until the last day of experiment (4 dpi). Because the objective of this work was to develop a protocol for isolating intact ECM from planarians, we did not characterize neither the proliferation rate nor the apoptotic event of the recellularized neoblasts. In any case, we observed no sign of nuclear fragmentation in the replanted cell, implying that there was little or no apoptotic event (Fig. 3K) (27). It would be of great interest to determine whether the neoblast could proliferate and inside the threedimensional microenvironment of the ECM-body. Further work should also further be conducted to monitoring the alteration in behaviour of repopulating cell. Optimization of the culture condition to promote other cellular processes such as migration, differentiation, stem cell renewal and eventually immortalization of the neoblast cells in the ECM-body would be greatly beneficial for the planarian community.

## Conclusions

We developed a simple workflow to isolate intact whole-body ECM from two highly potent freshwater planarians, *D. sp*. and *D. japonica*, using detergent-assisted decellularization. It is clear that the cell-free ECM retained the size and the overall shape of the worm. DAPI staining confirmed that the ECM-body isolated by the developed procedure has no cell contaminated. Scanning electron microscopic characterization revealed a uniformed porous structure on the surface of isolated ECM. Further characterization using Masson’s Trichrome histological staining clearly indicated faithful preservation of internal organization of the ECM-body. Taken together, the results thus validated the integrity of the structure of the isolated ECM. Finally, microinjection of FACS isolated X1 neoblasts into the ECM-body showed that not only the ECM-body could provide sites of attachment for the neoblast stem cells but also offer support for the survival of the cells. This work thus provides myriad of opportunities for planarian research specially to understand the effect of microenvironment in stem cell regeneration.

## Materials and Methods

### Animals

Freshwater planarian, *D. japonica*, were maintained in artificial spring water at room temperature (24 °C) and were fed twice a week with chicken liver. Animals were starved for at least 1 week before experiment.

### Isolation of planarian ECM-body

Planarians were terminated and stabilized in stabilization solution composed of 1% Nitric acid, 50 μM MgSO4 and 0.8% Formaldehyde. After that, the animals were decellularized in decellularization solution I composed of 0.08% SDS, EDTA pH 8.0 5mM, PMSF 1mM, and Tris-HCl pH 7.6 10 mM for 18 hours. Next, the solution was replaced with decellularization solution II composed of 0.04% SDS, EDTA pH 8.0 5mM, PMSF 1mM, and Tris-HCl pH 7.6 10 mM and incubated for at least 2 hours. Isolated ECM was then washed in wash solution composed of EDTA pH 8.0 5mM, PMSF 1mM, and Tris-HCl pH 7.6 10 mM for at least three times. ECM was finally stored at 4 °C in a freshly prepared washing solution. For the ECM aimed for cell culture, an additional nuclease treatment was added to remove potential contamination of small nucleic acid fragments. Afterward, samples were intensively washed with sterile PBS. Then, samples were cleaned with culture medium supplemented with antibiotics. Isolated ECMs were kept by submerging them in sterile PBS at 4 °C prior to use in the experiment. Detailed protocol is provided in the supplementary Materials and Methods.

### Characterization of decellularization efficiency using DAPI staining

DAPI staining was conducted to determine the presence of cells in the ECM-body. At first, the mucus layer of the worm was removed to enhance the accessibility of the solution. Both ECM-body and the worm were then fixed in formaldehyde solution. Afterward, the samples were incubated in permeabilizing solution. After that, the samples were incubated in DAPI solution for an hour in the dark. Stained samples were washed to remove the excess amount of DAPI and submerged in glycerol solution to increase the clarity of the sample. Mounted samples were observed under fluorescent microscope.

### Histological Characterization

Masson’s trichrome histological staining was conducted to evaluate the internal structure and general tissue architecture.

### Scanning Electron Microscopy

For ultrastructural investigation, both of samples were fixed with Glutaraldehyde and Osmium tetroxide. Planarians was dehydrated in ethanol whereas ECM-body was dehydrated in acetone. Subsequently, the samples were proceeded to critical point drying, platinum coating and mounted on SEM stub. The samples were visualized under scanning electron microscope.

### Fluorometric quantification of DNA and RNA in ECM

The amount of DNA and RNA in the ECM-body were evaluated by nanophotometer and fluorometer respectively.

### Planarian cell dissociation and neoblast microinjection into ECM-body

The preparation of single-cells for fluorescence-activated cell sorting analysis was conducted as described in Hayashi et al., 2006. X-ray sensitive neoblast cells (X1 population) were centrifuged and proceeded to injection. Aggregated cells were equally injected into posterior and anterior of pharynx and incubated in the culture medium at 24°C in CO2 incubator. After 1 and 3 days post injection, the ECM-body was stained with Hoechst 33342 or DAPI to visualize in injected cell and observed under confocal microscope (FV1000, Olympus). Detailed protocols of neoblast sorting and cell injection are stated in the supplementary Materials and Methods.

## Acknowledgement

The work was supported by the Center of Excellence on Environmental Health and Toxicology, Office of Higher Education Commission, Ministry of Education, Thailand and Thailand Research Funds (RSA5980078 to PO). We also thank financial support from Faculty of Science, Mahidol University. Oversea exchange scholarship from Gakushuin University and Mahidol University to ES is gratefully acknowledged. We thank S. Singhakaew for experimental assistance, S. Leetacheewa and K. Boonthaworn for helpful discussion.

## Author Contributions

PO and AC conceived the idea and designed the experiment. KA and TI further improved experimental design. ES, CW, MI and AM performed the experiments. SM and PW helped with characterization of decellularized ECM. KC and NC helped with SEM data collection. PO, TI and KA supervised the project. ES and PO wrote the original manuscript. All authors have reviewed and approved the manuscript.

## Supplementary Information

### Supplementary Materials and Methods

#### Isolation of Planarian Extracellular Matrix (ECM-body)

Seven-day starved planarians were collected from the tank. This protocol was designed for 40 planarians. Planarian was terminated and stabilized once at a time by replacing solution with stabilization solution (Ingredients of different solutions used for isolating planarians were listed in table S1) and incubated at 4 °C for 1 hour without shaking. After the stabilization, the solution was replaced with 40 ml of decellularization solution I for 16-18 hours at 4 °C. However, incubation time could be varied depending on the size and species of planarians. Subsequently, we replaced the solution with 40 ml decellularization solution II for 2 hours at 4 °C. After that, the solution was gently replaced with wash solution for 20 minutes 3 times in 4 °C on a seesaw rocker. After this incubation step, the prepared ECM-body are ready for characterization using various visualization techniques. In case of ECM-body aimed to prepare as a biomimetic scaffold for 3D cell culture, additional nuclease treatment step was added to remove residual nucleic acid fragments contaminated in the sample. The treatment was incubated for 1 hours at 37 °C without shaking. The isolated ECM-body was subjected to extensive washing in sterile 1X at 4 °C PBS for 5 times, 10 minutes each. The sample was agitated gently on a seesaw rocker. The solution was washed with LCDM and EPSCM (Yang et al., 2017) supplemented with antibiotic for at least 3 times for 10 minutes each and incubated at 24 °C on a seesaw rocker. Isolated ECM-body were maintained in sterile PBS at 4 °C prior to use in the experiment.

#### Nucleus Staining by DAPI and Hoechst 33342

Isolated extracellular matrix as well as intact animals were transferred into 24-well plate. A coverslip was added at the bottom of each well. The worm was treated with 5% N-Acetylcysteine for 10 minutes at room temperature to terminate the worm and wash away all the excess mucus. The samples were fixed in 0.8% formaldehyde (Sigma) for 1 hour at 4 °C. The samples were rinsed twice with PBSTx, or PBS supplemented with 0.3% Triton X-100 (Sigma). Samples were then incubated for 2 hours in a permeabilizing solution composed of 1X PBS and 0.5% Triton X-100 at 4 °C, and then rinsed thrice with PBSTx prior to stain with 1:400 DAPI solution (Invitrogen). The samples were then incubated in PBSTx in dark condition for 1 hour at room temperature. Stained specimen was then washed 5 times in PBSTx, for 5 minutes each. The samples were then subjected to incubating in 80% glycerol for 30 minutes at 4 °C. Finally, samples were transferred and mounted using 80% glycerol on a clean glass slide. DAPI stained sample was visualized under Olympus BX53 fluorescent microscope (Olympus, Singapore). For the samples stained with Hoechst 33342, samples were fixed with 4% paraformaldehyde, 5% methanol, PBS for 20 minutes, followed by double washing in PBSTx, for 10 minutes each. Subsequently, the solution was replaced with 1:2000 Hoechst 33342 in 1×PBS and incubated for 10 minutes. The samples were observed under Fluoview FV3000 confocal laser scanning microscope (Olympus).

#### Masson’s trichrome staining

Specimen was fixed in 1% Nitric acid, 50 μM MgSO4 and 0.8% formaldehyde overnight. Then, the ECM was dehydrated in the serial dilution of acetone (Fisher Scientific) while the dehydration of planarian was done in dilution series of ethanol. Dehydrated samples were further incubated in 1, 4-Dioxane (Fisher Scientific) for 30 minutes. The samples were embedded in the paraffin thrice for 20 minutes each. The paraffin block was trim at a thickness of 5 μm using ultramicrotome (Leica Biosystems). The sections were proceeded by a routine protocol for trichrome staining, which is good for staining muscle fiber, collagen, cytoplasm and nucleus staining.

#### Ultrastructure of planarian extracellular matrix

To prepare sample for observation under scanning electron microscope, ECM of planarian were collected and fixed with 2.5% glutaraldehyde in 0.1 M sodium cacodylate pH 7.2 at 4 °C and washed with 0.1 M sodium cacodylate buffer pH 7.2 at 4 °C. Next, ECM were fixed with 1% osmium tetroxide in 0.1 M sodium cacodylate buffer for 1 hour at 4 °C. After that, the samples were washed with 0.1 M sodium cacodylate buffer pH 7.0 at 4 °C. Worms were dehydrated by soaking in series of concentration of ethyl alcohol which were 30%, 50%, 70% 80%, 90% and 95% for 15 minutes each, at 4 °C. After the samples are in 95% ethyl alcohol, absolute ethyl alcohol was used to replace the solution at 4 °C. The residual ethyl alcohol was removed using critical point drying (CPD). The samples were mounted on stubs with double-coated conductive carbon tape. Specimen was coated with Pt-Pd for 4 minutes using ion sputter before observing under JSM-6510 Series Scanning Electron Microscope (JEOL).

#### Fluorometric Quantification of DNA and RNA in Extracellular Matrix

Nucleic acid material remained in extracellular matrix were quantified using fluorometric method. Commercial kit (Bio Basic Canada Inc.) was utilized to extract DNA from the ECM sample in comparison with the intact planarian. The concentration of DNA was determined by nanophotometer (Implen, Germany). For RNA detection, RNA was extracted all ECM and intact planarian using Trizol reagent (Invitrogen). After extraction, the RNA was resuspended in mild base water. The concentration of RNA was determined by QFX Fluorometer (DeNovix).

#### Planarian Cell Dissociation and Neoblast Microinjection

Planarian was cut into 3-4 fragments by clean scalpel. The fragments were then soaked in Holtfreter’s solution diluted in distilled water. The fragments were further subjected to physical mincing and treatment with 0.25% trypsin for several minutes at 20 °C. The cells were dissociated by gentle pipetting up and down several times. The dissociated cells were orderly filtered through a 35 μm pore size cell strainer and 20 μm nylon net filter to remove tissue fragments. Single-cells suspension was stained with Dye Cycle (Invitrogen) for 30 minutes at room temperature. Flow cytometric analysis was performed using a BD FACS Melody Cell Sorter (Becton-Dickinson). X-ray-sensitive neoblast cell or X1 fraction were isolated and utilized in the microinjection. 1×105 X1 cells could be successfully temporarily maintained in 24-well plate in 1 ml culture medium on a rotary shaker at 20 0C in CO2 incubator. The cells were collected by centrifuge at 300xg for 7 minutes. 10 μl of aggregated cells were drawn into capillary glass of the injector. ECM-body was placed firmly on the black sterile filter paper for the injection. The cells were then injected into the posterior and anterior parts of the pharynx. After the injection, ECM-body was transferred gently into culture medium and cultivate at 24°C in a CO2 incubator.

#### SDS-PAGE analysis of ECM components

Planarian was cut into 3-4 fragments by clean scalpel. The fragments were then soaked in Holtfreter’s solution diluted in distilled water. The fragments were further subjected to physical mincing and treatment with 0.25% trypsin for several minutes at 20°C.

### Supplementary Table

**Table S1:**
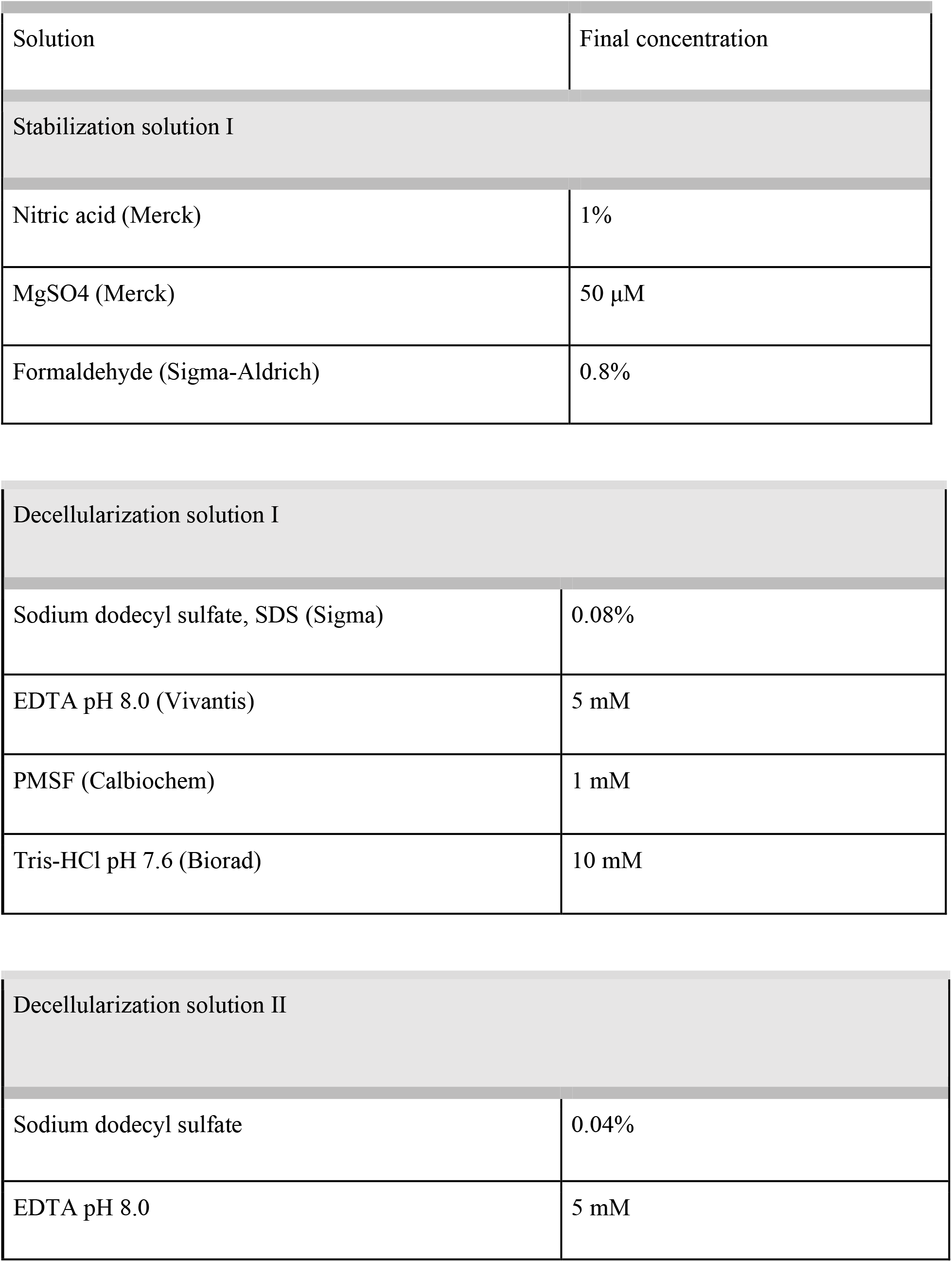
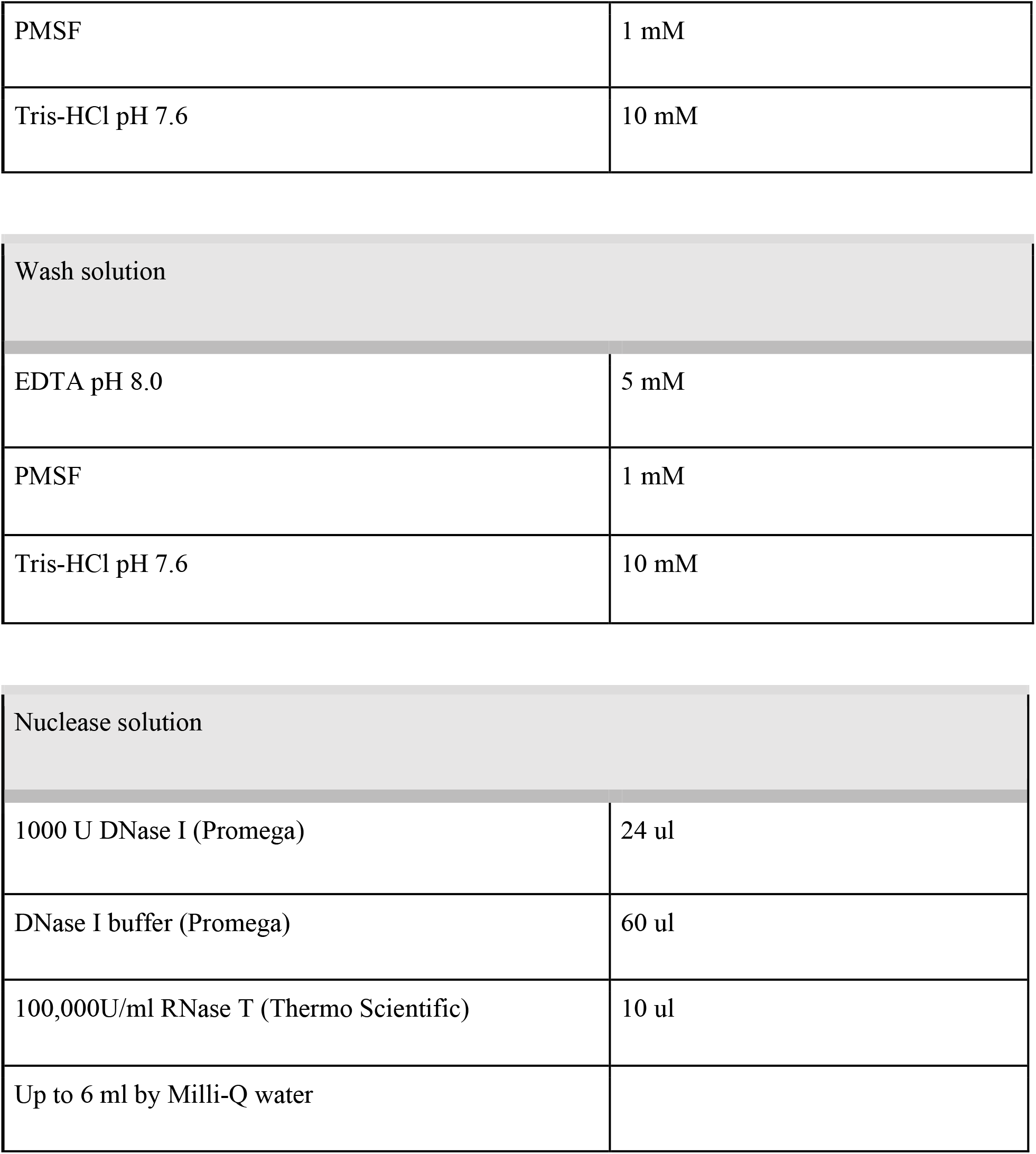
Buffer Recipes used in Planarian Decellularization

| Solution | Final concentration |
| --- | --- |
| Stabilization solution I |  |
| Nitric acid (Merck) | 1% |
| MgSO<sub>4</sub> (Merck) | 50 μM |
| Formaldehyde (Sigma-Aldrich) | 0.8% |
| Decellularization solution I |  |
| Sodium dodecyl sulfate, SDS (Sigma) | 0.08% |
| EDTA pH 8.0 (Vivantis) | 5 mM |
| PMSF (Calbiochem) | 1 mM |
| Tris-HCl pH 7.6 (Biorad) | 10 mM |
| Decellularization solution II |  |
| Sodium dodecyl sulfate | 0.04% |
| EDTA pH 8.0 | 5 mM |
| PMSF | 1 mM |
| Tris-HCl pH 7.6 | 10 mM |
| Wash solution |  |
| EDTA pH 8.0 | 5 mM |
| PMSF | 1 mM |
| Tris-HCl pH 7.6 | 10 mM |
| Nuclease solution |  |
| 1000 U DNase I (Promega) | 24 ul |
| DNase I buffer (Promega) | 60 ul |
| 100,000U/ml RNase T (Thermo Scientific) | 10 ul |
| Up to 6 ml by Milli-Q water |  |

